# Antagonism by the Type VI secretion system of *Bacteroides fragilis* is controlled by a TetR family regulator and released small molecule

**DOI:** 10.1101/2025.08.20.671311

**Authors:** Leila Tuzlak, Téa E. Pappas, Michael J. Coyne, Madeline L. Sheahan, Victoria Burgo, Phoebe A. Rice, Laurie E. Comstock

**Affiliations:** Duchossois Family Institute University of Chicago; Chicago, IL; Department of Microbiology, University of Chicago; Chicago, IL; Committee on Microbiology, University of Chicago, Chicago, IL; Department of Biochemistry and Molecular Biology, University of Chicago, Chicago. IL

**Keywords:** antagonism, microbiota, *Bacteroides*, T6SS, TetR

## Abstract

Antagonistic systems of bacteria are often tightly regulated. The human gut Bacteroidales harbor three distinct antagonistic Type VI secretion systems (T6SS), one of which is present only in *Bacteroides fragilis*, known as the GA3 T6SS. Although this is the best studied of the three T6SSs, little is known about how it is regulated. The gene upstream of the GA3 T6SS locus encodes a TetR family transcriptional regulator (TetR_GA3_), which we show represses expression of the GA3 T6SS locus. The gene immediately upstream and divergently transcribed from *tetR*_GA3_, designated here as *lgs*_GA3_, encodes a product of the α-oxoamine synthase family of pyridoxal phosphate-dependent enzymes with structural homology to the CqsA autoinducer synthase of the CAI-1 quorum sensing system of *Vibrio spp*. When *lgs*_GA3_ is deleted, transcription of the GA3 T6SS locus is repressed in a TetR-dependent manner. Strains synthesizing Lgs_GA3_ produce a molecule present in the supernatant that likely serves as the TetR_GA3_ ligand, overcoming TetR transcriptional repression of the GA3 T6SS. We show that GA3 T6SS-specific immunity genes present on two acquired immunity defense islands are also regulated by Lgs_GA3_ coordinating expression of GA3 T6SS antagonism with protection from competitor’s GA3 T6SS toxins. Production and firing of the GA3 T6SS and subsequent antagonism occurs in bacteria deleted for *lgs*_GA3_ when grown with bacteria containing this gene or their supernatants. These data show that the GA3 T6SS is regulated by a small molecule acting through TetR_GA3_ allowing the bacteria to coordinate antagonistic and protective systems.

**Significance statement:** There are numerous external and intrinsic signals that dictate when bacteria become aggressive and when they activate their defensive systems. We show that *B. fragilis* strains with a GA3 T6SS synthesize a small molecule released from bacterial cells that acts through TetR family regulators to coordinate transcription of both the antagonistic GA3 T6SS and arrays of immunity genes to competitor’s GA3 T6SS toxins. Bacteria can respond to this molecule when released from non-kin bacteria, allowing them to sense and respond to a threat from a nearby competitor. The coordinated regulation of the GA3 T6SS and arrays of immunity genes is the second example of regulatory crosstalk between the GA3 T6SS and genes of MGEs through TetR family regulators.

## Introduction

Bacteria that live in communities participate in numerous interactions, including antagonizing competing strains and species. Toxins produced by antagonistic systems can be deployed by various mechanisms including secretion of diffusible toxins that bind receptors on target cells and the direct injection of toxins into target cells by dedicated toxin delivery secretion systems such as the Type VI secretion system (T6SS) (reviewed (1, 2)). The deployment of antibacterial toxins is a tightly regulated process in many bacteria (3-7). Bacterial members of the human gut microbiota engage in extensive antagonistic behaviors (3). Bacteroidales is an order of bacteria with numerous species of the human gut microbiota that are collectively the most abundant order of Gram-negative bacteria in this ecosystem (8-10). These bacteria deploy numerous diffusible toxins (11-18), and have three distinct T6SSs known as GA1, GA2, and GA3 (19).

T6SSs are nanomachines built on the inside of the bacterial cell with a toxin-laden tube wrapped in a sheath that contracts to inject its toxins into neighboring cells. In many bacteria, synthesis and firing of the apparatus is tightly controlled to limit firing until a threat is detected (4, 20). We know little about how the T6SSs of the gut Bacteroidales are regulated *in vitro* or in the mammalian gut. We previously showed that each of the three T6SSs of gut Bacteroidales has an adjacent gene of the TetR family of transcriptional regulators (19). Conjugal acquisition of the ICE containing the GA1 T6SS by GA3 T6SS-containing *Bacteroides fragilis* leads to transcriptional repression of the GA3 T6SS locus by the TetR encoded adjacent to the GA1 T6SS locus (TetR_GA1_) (21).

TetR family regulators comprise one of the largest families of bacterial transcriptional regulators (reviewed (22-24)). The importance of these proteins in regulating antibiotic resistance, central metabolism, biofilm formation, virulence, quorum sensing and numerous other properties has been well documented (reviewed (22)). Despite sharing little primary sequence similarity, TetR family regulators each have an N-terminal DNA binding domain and a C-terminal ligand binding domain. TetR family regulators typically bind as a homodimer to a palindromic DNA sequence near the promoter region of target genes, repressing transcription until a specific ligand binds the C-terminal region. Ligand binding releases TetR and thus allows transcription of the downstream genes. The ligands that bind and de-repress transcription have yet to be identified for many well-characterized TetR family proteins (23).

Here, we set out to elucidate the involvement of the TetR encoded directly upstream of the GA3 T6SS locus (TetR_GA3_) in regulation of its downstream region. We show not only that TetR_GA3_ represses expression of the GA3 locus, but that the gene upstream of *tetR*_GA3_ encodes an enzyme that synthesizes a small molecule that overcomes this repression. We show that this small molecule is released from the bacteria and that supernatants from non-kin bacteria producing this enzyme can overcome TetR_GA3_ repression of the GA3 T6SS and of acquired arrays of immunity genes to heterologous GA3 T6SS toxins.

## Results

The gene immediately upstream of the GA3 T6SS locus encodes a TetR family regulator designated TetR_GA3_ (Fig 1a). To determine if TetR_GA3_ regulates its downstream T6SS locus, we overexpressed *tetR*_GA3_ either in single copy from a constitutive promoter or from a multi-copy plasmid with the same constitutive promoter and assayed for production of the most abundant GA3 T6SS component, the main structural tube protein of the GA3 T6SS (25), previously referred to as Hcp1 and herein referred to as Hcp_GA3_. Analyses of stationary phase cultures using two different *B. fragilis* strains, 638R and NCTC9343 (9343), revealed that the single additional copy of *tetR*_GA3_ does not result in obvious changes in Hcp_GA3_ levels compared to WT in either cells or the supernatant (a proxy for T6SS firing). However, TetR_GA3_ production from the multicopy plasmid completely inhibits synthesis of Hcp_GA3_ (Fig 1b). To elucidate the TetR_GA3_ regulon, we performed transcriptomic analysis of strain 638R with the multi-copy *tetR*_GA3_-containing plasmid compared to the empty vector control strain. These data show that TetR_GA3_ regulates its own transcription, the GA3 T6SS locus, the gene immediately upstream and divergently transcribed from *tetR*_GA3_ (BF638R_1996), as well as two other *tetR* containing loci, drastically reducing expression of these genes (Fig 1c, 1d, 1e, Dataset 1, tab 1). These two other loci are part of mobile genetic elements (MGE) known as acquired interbacterial defense (AID) systems (26). These AID each contain genes encoding TetR orthologs that are very similar to TetR_GA3_ (Fig S1B) and arrays of immunity genes to GA3 T6SS effectors previously described as *B. fragilis* specific orphan immunity gene cluster 2 and AID1 (cluster 1) (26) (Fig 1e).

**Figure 1.**
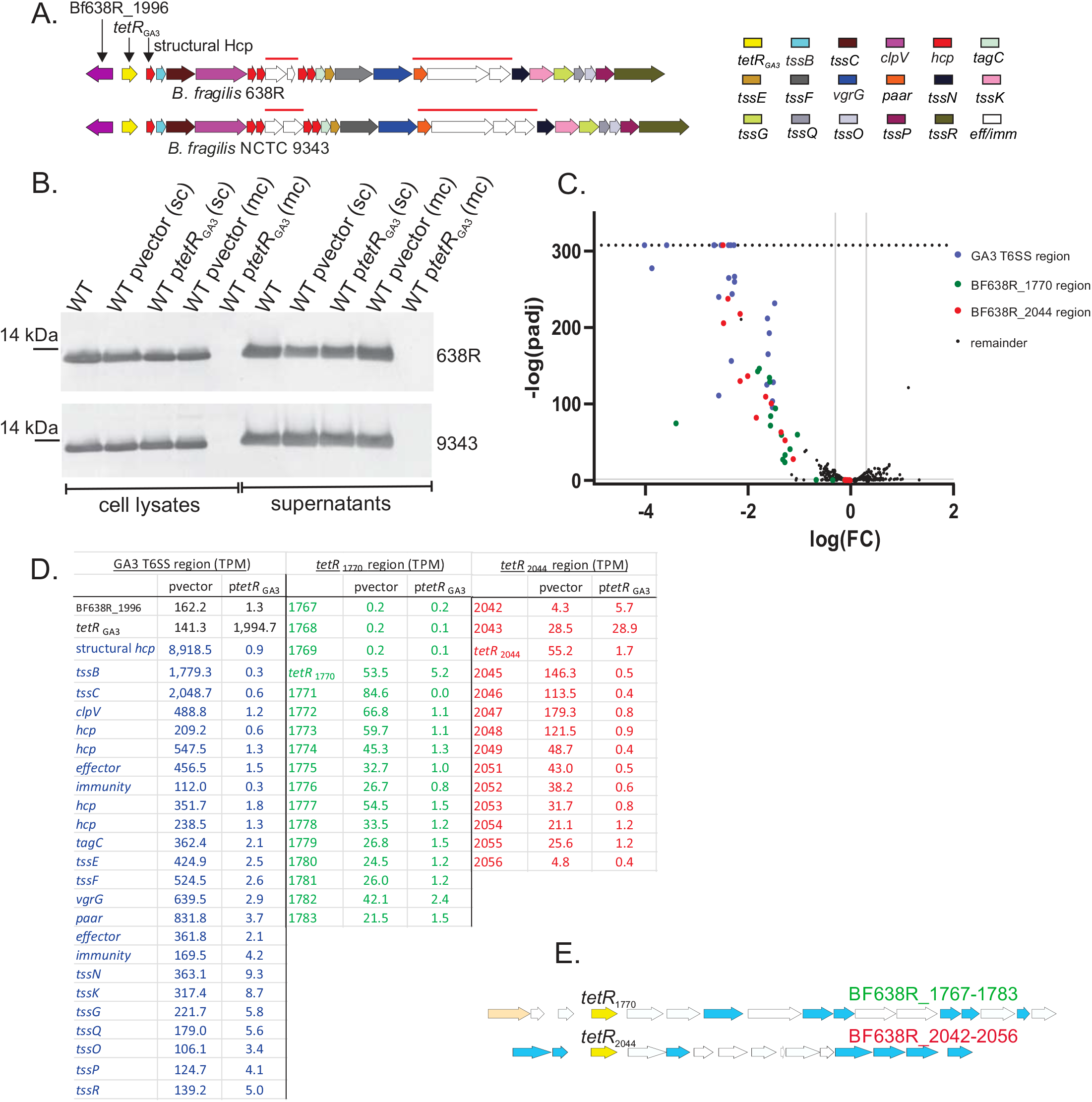
TetR_GA3_ regulates expression of the GA3 T6SS locus and two acquired interbacterial defense loci. **A**. Gene maps of the T6SS loci and two conserved upstream genes from two *B. fragilis* strains. Red lines over genes indicate two variable regions encoding toxic effector and immunity genes. **B**. Western immunoblot of cell lysates and supernatants from strains 638R and 9343 expressing *tetR*_GA3_ from a single copy (sc) integrated construct or from a multicopy (mc) replicating plasmid. Blots were probed with antiserum to the structural Hcp protein (Hcp_GA3_) encoded by the first gene of the locus (BF638R_1994). Loading control blots shown in Fig S1A. **C**. Volcano plot of transcriptomics data comparing strain 638R overexpressing *tetR*_GA3_ from the multicopy plasmid to the vector control strain. In instances where both the padj (from DESeq2) and the FDR (from EdgeR) values were zero, these values were replaced with 2.22507385850721E-308, the smallest positive number Excel can utilize, to allow calculation of the corresponding negative log value (horizontal line). Genes corresponding to the GA3 T6SS locus and the two immunity defense islands are colored. **D**. Transcripts Per Million (TPM) values are shown for the three regions that are drastically repressed when *tetR*_GA3_ is overexpressed. **E**. Gene maps of the two acquired interbacterial defense islands that are repressed by TetR_GA3_. A *tetR* gene in each locus is colored yellow and GA3 T6SS immunity genes identified in (26) are colored light blue.

Although these immunity genes are specific to counteract GA3 T6SS effectors, the immunity genes of these two regions do not counteract effectors of the GA3 T6SS of strain 638R, but rather incoming toxic effector from competitors (26).

The ligands of TetR family regulators are diverse and are often exogenous molecules such as antibiotics, bile acids, or other toxic molecules (23). The fact that Hcp_GA3_ synthesis is repressed by TetR_GA3_, yet Hcp_GA3_ is abundant in both the cellular and supernatant fractions of broth-grown cultures suggests that the TetR_GA3_ ligand may be a bacterial product. The gene immediately upstream of *tetR*_GA3_, (BF638R_1996) is drastically downregulated when *tetR*_GA3_ is overexpressed (Fig 1D), and we previously showed it is also drastically downregulated by TetR_GA1_ (21). This gene encodes a product that belongs to the α-oxoamine synthase family of PLP-dependent enzymes. These enzymes perform a condensation reaction between an amino acid substrate and acyl-CoA and include the enzymes 5-aminolevulinate synthase, 8-amino-7-oxononanoate synthase, and serine palmitoyl transferase. The BF638R_1996 enzyme is also predicted to be structurally related to CqsA (Fig 2a), of *Vibrio spp*. that synthesizes the autoinducer precursor molecule amino-CAI-1 (27, 28). Interestingly, a closely related protein is encoded between two *tetR* orthologs adjacent to the Bacteroidetocin-B locus, a four-gene region that produces the bacteriocin Bd-B (14) (Fig 2b). This intervening gene (EH213_01840) encodes a product that is 78% similar to BF638R_1996 (Fig 2a, 2b, Fig S2C). 3D similarity analysis using the AlphaFold-predicted structures from these products, BF638R-1996 and EH213_01840, showed a root mean square deviation (rmsd) of 0.84Å for 416 structurally aligned Calpha atoms (Fig 2a), whereas alignments of these structures to the CqsA experimental structure gave rmsds of 2.58Å and 2.61Å, respectively, over 355 Calpha atoms. As a control, the AlphaFold-predicted structure of CqsA could be superimposed on the experimental structure with an rmsd of 0.58Å over 386 Calpha atoms. These data show that genes encoding very similar enzymes are linked to *tetR* family genes of loci encoding toxin/antagonistic functions. Based on their structural similarity, these two putative α-oxoamine synthase PLP-dependent enzymes are likely to produce similar small molecules that we predicted are the ligands for their respective TetR regulators.

**Figure 2.**
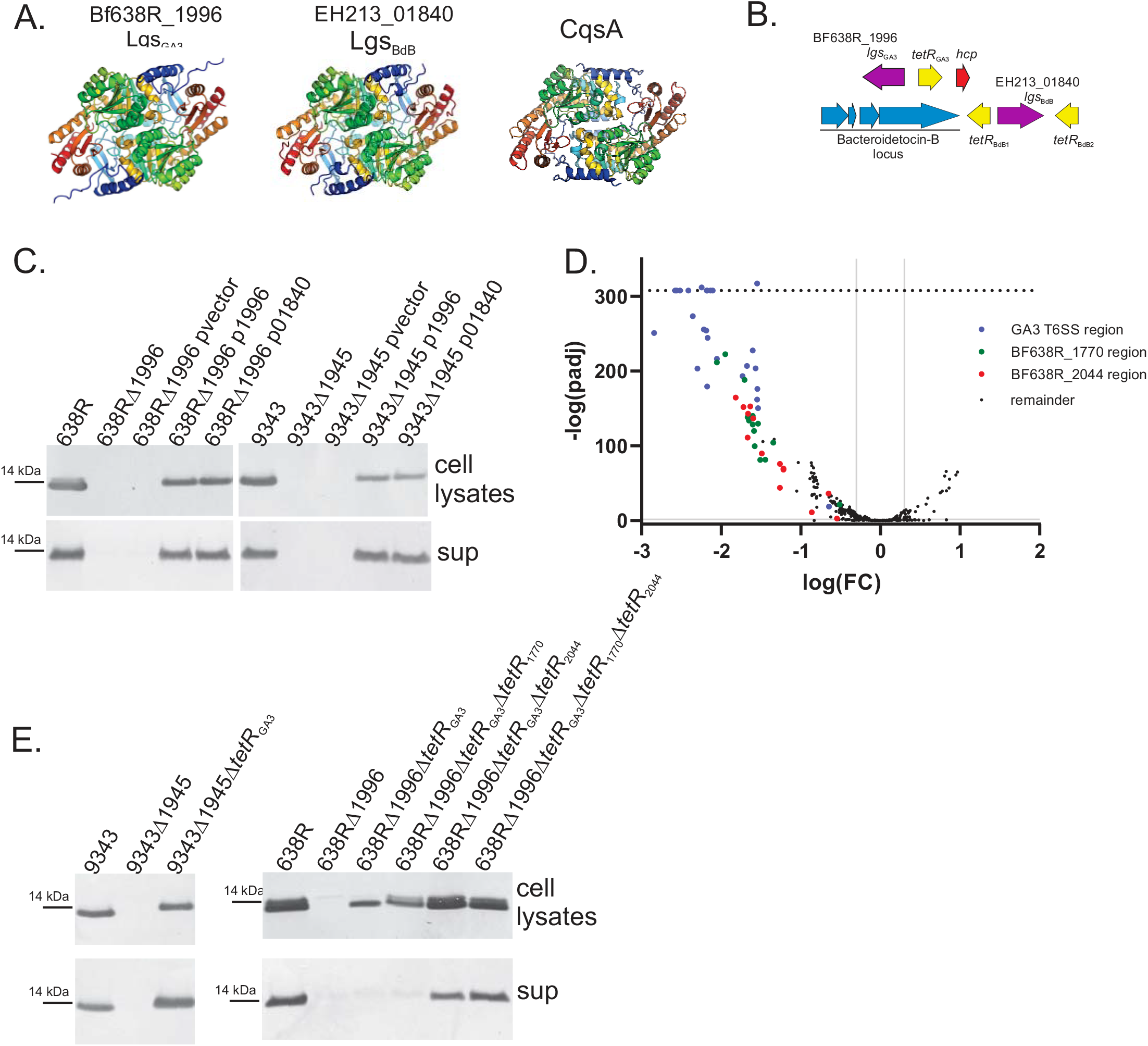
The gene upstream of *tetR*_GA3_ encodes an enzyme required to overcome TetR_GA3_ repression. **A**. AlphaFold3-predicted dimer structure of enzymes encoded by BF638R_1996 and EH213_01840 and the apo CqsA dimer structure (PDB ID 2wk7) (27). Each protein chain is shaded from blue (N-ter) to red (C-ter). Inter- and intra-molecular scores for the BF638R_1996 model: ipTM = 0.82 pTM = 0.84 and the E213_01840 model: ipTM = 0.8 pTM = 0.85. Figure prepared with PyMOL (https://pymol.org/). **B**. Gene maps showing the *tetR* and surrounding genes of the GA3 T6SS and bacteroidetocin-B region of strain *B. thetaiotaomicron* CL15T12C11. **C**. Western immunoblot of cell lysates and supernatants of strains 638R and 9343 deleted for the gene encoded immediately upstream of *tetR*_GA3_ (BF638R_1996 or BF9343_1945), and these mutants strains complemented with the native gene or the gene from the Bd-B region (01840). Blots were probed with antiserum to Hcp_GA3_. Loading control blots shown in Fig S2A. **D**. Volcano plot of the transcriptomics data comparing strains 638R and 638RΔ1996 (Δ*lgs*_GA3_). Genes corresponding to the GA3 T6SS locus and the two immunity defense islands are colored.In instances where both the padj (from DESeq2) and the FDR (from EdgeR) values were zero, these values were replaced with 2.22507385850721E-308, the smallest positive number Excel can utilize, to allow calculation of the corresponding negative log value (horizontal line). **E**. Western immunoblot of cell lysates and supernatants of strains 638R and 9343 deleted for BF638R_1996 or BF9343_1945 and mutants with deletions of these genes and their adjacent *tetR*_GA3_ gene. For strain 638R, deletions of the *tetR*_1770_ and *tetR*_2044_ were also made in the Δ1996Δ*tetR*_GA3_ background. Blots were probed with antiserum to Hcp_GA3_. Loading control blots are shown in Fig S2B.

To determine if the *B. fragilis* gene encodes a product involved in the regulation of the GA3 T6SS locus, we deleted it from strains 638R (gene BF638R_1996) and 9343 (gene BF9343_1945). Western immunoblot analysis showed complete abrogation of Hcp_GA3_ synthesis in both these mutants (Fig 2C). Hcp_GA3_ production was restored when the gene was reintroduced to the mutant strains, and surprisingly, the ortholog from the Bd-B locus (EH213_01840) similarly restored Hcp_GA3_ synthesis (Fig 2C), providing further evidence that these related enzymes produce the same or functionally similar molecule. Transcriptomic analysis of the BF638R_1996 deletion mutant revealed an expression profile similar to the 638R strain overexpressing t*etR*_GA3_, with the GA3 T6SS and the two immunity gene regions drastically downregulated (Fig 2D, Dataset 1 tab 2). If the Bf638R_1996 enzyme produces the molecule that serves as the TetR_GA3_ ligand, it would be expected that further deletion of *tetR*_GA3_ from these mutant strains would restore Hcp_GA3_ synthesis. Indeed, deletion of both genes together resulted in complete restoration of Hcp_GA3_ synthesis in strain 9343 (Fig 2E); however, Hcp_GA3_ production was only partially restored in 638R (Fig 2E). Strain 638R produces TetR_1770_ and TetR_2044_ of the immunity gene MGEs, which are both similar to TetR_GA3_ (Fig S1B), whereas strain 9343 does not contain any TetR orthologs similar to TetR_GA3_ that may inhibit GA3 T6SS production. We found that when *tetR*_2044_, but not *tetR*_1770_ is deleted from 638RΔ1996Δ*tetR*_GA3_, levels of Hcp_GA3_ are restored to near wild-type level (Fig 2E). These data show that the gene upstream of *tetR*_GA3_ encodes an enzyme that produces a molecule necessary to de-repress TetR_GA3_ and TetR_2044_ inhibition of Hcp_GA3_ synthesis. These data strongly suggest that the BF638R_1996 enzyme produces the TetR_GA3_, TetR_2044_ and likely TetR_1770_ ligand, and we therefore designated it Lgs_GA3_.for ligand synthase GA3.

We performed two analyses to determine if the Lgs_GA3_ product is released from bacterial cells and present in culture supernatants, similar to amino-CAI-1. First, we co-cultured 638RΔ*lgs*_GA3_ with various strains producing *lgs*_GA3_ or the orthologous gene of the Bd-B locus, *lgs*_BdB_. The strains used for co-culture included 638RΔGA3 (deleted for the entire GA3 T6SS locus but retaining *tetR*_GA3_ and *lgs*_GA3_), *Bacteroides thetaiotaomicron* CL15T12C11ΔBdB, deleted for the bacteriodetocin-B gene so that it cannot kill 638R (14), BtCL15T12C11ΔBdBΔ*lgs*_BdB_, and *B. thetaiotaomicron* VPI-5482 with an empty vector, or with the multi-copy plasmid expressing *lgs*_GA3_ or *lgs*_BdB_. *B. thetaiotaomicron* VPI5482 does not have a T6SS or Bd-B locus or orthologs with high similarity to Lgs_GA3_. In all cultures where 638RΔ*lgs*_GA3_ was co-cultured with a strain expressing either *lgs*_GA3_ or *lgs*_BdB_, Hcp_GA3_ was produced at high levels and the GA3 T6SS of 638RΔ*lgs*_GA3_ fired as evidenced by Hcp_GA3_ in the supernatant (Fig 3A). Therefore, transfer of a single gene, *lgs*_GA3_ or *lgs*_BdB_, to a heterologous strain that does not contain any of this antagonistic or regulatory machinery, is sufficient to produce a molecule that is recognized by the 638RΔ*lgs*_GA3_ strain to overcome TetR_GA3_ repression to produce and fire its GA3 T6SS. As further evidence for the extracellular presence of the ligand, we used filtered culture supernatants from the same strains used for co-culture and showed that although Hcp_GA3_ production is less due to a finite amount of ligand in these supernatants, they restored Hcp_GA3_ synthesis to the 638RΔ*lgs*_GA3_ mutant (Fig 3B, top panel). To further show that the ligand acts by depressing the inhibitory activity of the TetR_GA3_, we performed the same growth experiment using cell supernatants from Lgs-producing bacteria, but this time showed that they could derepress Hcp_GA3_ synthesis in the strain overexpressing *tetR*_GA3_ from the multicopy plasmid (Fig 3B, bottom panel). As we showed that TetR_GA3_ represses the ligand synthase gene when overexpressed from this vector, de-repression of Hcp_GA3_ synthesis in this strain could not occur unless exogenous ligand is added or produced from co-cultured strains.

**Figure 3.**
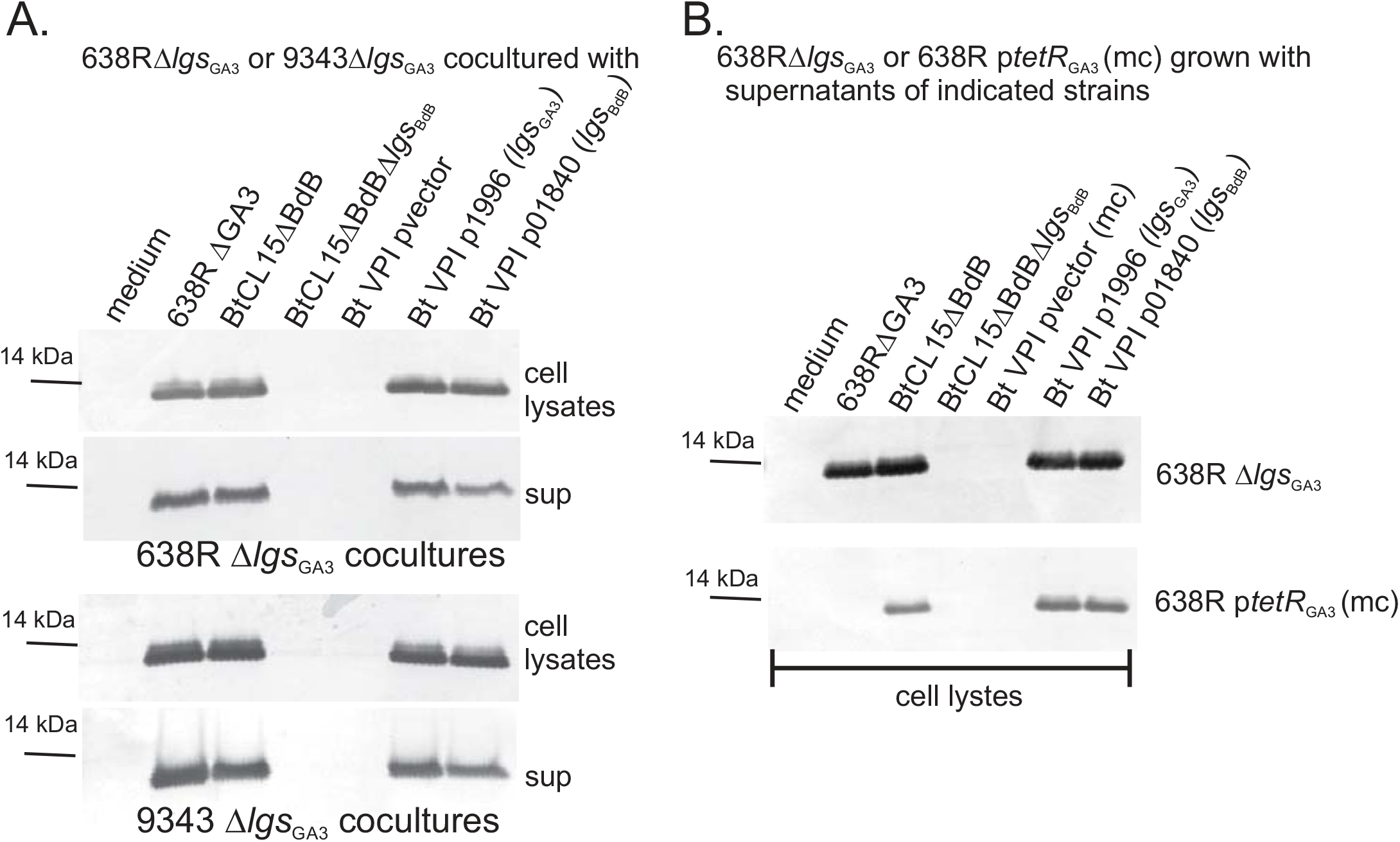
The molecule(s) produced by Lgs_GA3_ and Lgs_BdB_ are present in the cell supernatant and act on non-kin cells. **A**. Western immunoblots showing production of Hcp_GA3_ from 638RΔ*lgs*_GA3_ or 9343Δ*lgs*_GA3_ co-cultured with strains producing Lgs_GA3_ or Lgs_BdB_ compared to co-culture with control strains. Western immunoblot of loading controls are shown in Fig S3A. **B**. Western immunoblots showing production of Hcp_GA3_ from 638RΔ*lgs*_GA3_ or 638R p*tetR*_GA3_ (mc) grown with 50% filtered culture supernatants from strains producing Lgs_GA3_ or Lgs_BdB_ compared to supernatants from control strains. Fig S3B shows the western immunoblot of the supernatants added to the growth medium, showing they do not contain Hcp_GA3_. Western immunoblots of the loading controls are shown in Fig S3C.

These data suggest that through this form of regulation, i.e. sensing a molecule that likely interacts with TetR_GA3_ thereby derepressing GA3 T6SS and immunity gene expression, this system allows these bacteria to sense and turn on GA3 T6SS antagonism (and protection) when they are near other GA3 T6SS, or Bd-B-producing competitors. We performed antagonism assays to determine whether *B. fragilis* can sense the ligand produced by a cocultured strain to kill the competitor using the GA3 T6SS. As antagonistic and control strains, we used 638R, the 638R ΔT6 mutant that cannot fire the weapon (25), and 638R Δ*lgs*_GA3_, that does not produce the T6SS weapon in monoculture but can sense the ligand produced by Lgs_GA3_ and Lgs_BdB_ released by co-cultured bacteria. As targets, we used the 638R erythromycin resistant (*ermG*) isogenic strain in which the two variable regions of the GA3 T6SS locus encoding the toxic effector and immunity proteins (638R*erm*ΔV1ΔV2) are deleted (25), and therefore cannot antagonize or protect itself from GA3 T6SS antagonism. In addition, we deleted *lgs*_GA3_ from this 638R*erm*ΔV1ΔV2 strain and used it as an additional target strain. As the concentration of the ligand should be important for de-repression by TetR_GA3_, we used two different ratios of antagonist to target (5:1 and 1:1). Higher ratios of antagonist to target result in greater antagonism. At the 5:1 ratio, the wild-type 638R strain was able to antagonize to approximately 3-log when compared to the 638R ΔT6 non-antagonistic mutant. At this ratio, although the ligand was only being provided by ∼17% of cells in the initial co-culture (the target cells), there was still a 1-log killing by the 638R Δ*lgs*_GA3_ strain, only when co-cultured with the target strain producing Lgs_GA3_, but not by the target strain deleted for *lgs*_GA3.,_ At the 1:1 ratio, antagonism by the WT strain was reduced to 1-log, but the 638R Δ*lgs*_GA3_ strain matched that level of antagonism, as now 50% of the cells of the initial co-culture were producing the ligand, which appears adequate for WT levels of killing in this assay. These data show that this mechanism of regulation of GA3 T6SS antagonism allows bacteria to respond to ligand-producing competitors to initiate antagonism.

## Discussion

In this study, we sought to elucidate the role of TetR_GA3_ in the regulation of the GA3 T6SS of *B. fragilis*. Using two *B. fragilis* strains, we demonstrated that overexpression of TetR_GA3_ represses the GA3 T6SS locus, and that an enzyme (Lgs_GA3_) encoded by a gene adjacent to *tetR*_GA3_ produces a small molecule that is necessary to overcome TetR_GA3_ transcriptional repression of the GA3 T6SS locus.

Lgs_GA3_ and Lgs_BdB_ belong to the α-oxoamine synthase (AOS) family of PLP-dependent enzymes that perform a condensation reaction between an amino acid substrate and acyl-CoA. This large enzyme family includes aminolevulinate synthase involved in heme production (29), serine palmitoyltransferase, involved in sphingolipid synthesis (30) and CqsA of *Vibrio spp*. (31), which produces amino-CAI-1 (27, 28), that is converted into the quorum sensing autoinducer CAI-1 (3-hydroxytridecane-4-one) (32). AOS enzymes share a common conserved structural fold and have similar residues in their active sites (33). Based on the substrates of AOS enzymes, we predict that Lgs_GA3_ will similarly transfer an acyl group from acyl-CoA to an amino acid type molecule. We predict that the Lgs_GA3_ product is the TetR_GA3_ ligand, which binds the C-termini of the TetR_GA3_ dimer inducing a conformational change affecting its ability to bind target DNA. The products of Lgs_GA3_ and Lgs_BdB_ are likely very similar or identical as they both relieve transcriptional repression by TetR_GA3_. Further work is necessary to understand if and how the Lgs_BdB_ and the two TetR_BdB_ family repressors regulate the Bacteroidetocin-B locus (14).

Our studies show that the small molecule(s) produced by Lgs_GA3_ and Lgs_BdB_ enzymes are present in the extracellular milieu which may result from active transport, diffusion across membranes, and/or facilitated by outer membrane vesicle release as shown for CAI-1 (34). If the small molecule or a modified version is the TetR_GA3_ ligand as we predict, it likely diffuses across membranes reaching an equilibrium inside and outside the cell.

The product of Lgs_GA3_ also affects the inhibitory activity of TetR_1770_ and TetR_2044_ of the Bacteroidales acquired orphan immunity defense islands, AID-1 and AID-2 (26). These MGE do not appear to encode their own *lgs* homolog, suggesting that depression by these TetR relies on the Lgs_GA3_ enzymatic product, thereby coordinating antagonism with protection. In addition, a Bacteroidales strain lacking a GA3 T6SS but that contains these AID should sense and respond to the Lgs_GA3_ ligand released from a nearby GA3 T6SS-antagonizing *B. fragilis* strain to bind their AID TetR orthologs and turn on these arrays of protective genes. Future studies focused on identifying the structure of the Lgs_GA3_ product, how it interacts with TetR_GA3_, TetR_1770_, TetR_2044_, and identification of the TetR DNA binding motifs of all three loci will shed light on the precise molecular details of this regulatory system.

With our current knowledge of TetR_GA3_ and Lgs_GA3_ mediated regulation of the GA3 T6SS, we can now delve into the precise mechanism of how TetR_GA1_ encoded by the GA1 ICE, which is present in ∼18% of *B. fragilis* strains with a GA3 T6SS, inhibits transcription of the GA3 T6SS locus as previously described (21). *tetR*_GA1_ is not present near any genes that encode an enzyme with similar activity to Lgs_GA3_. Our previous data showed that *lgs*_GA3_ is transcriptionally inhibited by TetR_GA1_ and it is possible that this inhibition allows constitutive repression of the GA3 T6SS locus by TetR_GA3_. Alternatively, TetR_GA1_ may inhibit the GA3 T6SS in a manner independent of the activities of TetR_GA3_ and Lgs_GA3_. Interestingly, the GA2 T6SS locus is adjacent to two genes encoding TetR family regulators. Between these two *tetR*_GA2_ is a gene encoding a product with structural similarity to ketoacyl-synthase type enzymes (reviewed (35)). It is likely that one of the TetR_GA2_ proteins bind the small molecule produced by this keto-acyl-like enzyme. Therefore, we predict that regulatory networks involving TetR family regulators and adjacent small molecule synthesis genes control numerous antagonistic and protective systems of the gut Bacteroidales.

## Methods

All primers are listed in Table S1 and strains and plasmids are listed in Table S2.

### Strains and growth conditions

*Bacteroides* strains were grown under anaerobic conditions on BHIS plates (36), in basal liquid medium (37), or in modified basal medium (MBM), modified to prevent a sharp decline in pH at stationary phase (Leonor García-Bayona, personal communication). MBM is basal medium modified by adjusting the pH to 8.5 with NaOH prior to autoclaving. Prior to use, 5 ml of 1 M sodium bicarbonate was added to 100 ml basal along with the other supplements. Antibiotics used for selection or induction include erythromycin (10 µg/mL), gentamicin (200 µg/mL), cefoxitin (10 µg/mL), carbenicillin (100 µg/mL) and anhydrotetracycline (75 ng/mL).

### General methods for genetic constructs

DNA regions for all constructs were PCR amplified using Phusion or Q5 polymerase (NEB) with primers listed in Table S1. NEBuilder (New England Biolabs, Ipswich, MA) was used for all cloning and assembly of DNA segments into the BamHI site of vectors (indicated in Table S1). All recombinant plasmids were sequenced by Plasmidsaurus (Eugene, OR).

### Construction of deletion mutants

All deletion mutants were constructed by cloning the flanking segments of the region to be deleted into the BamHI site of pMLS36 (21) or pLBG13 (36) and transformed into *E. coli* S17-1 λ pir. Plasmids were conjugally transferred from *E. coli* to *Bacteroides* and co-integrates were selected on BHIS containing gentamicin (200µg/ml) with either cefoxitin (10µg/ml) or erythromycin (10 µg/ml). Cointegrates were passaged in basal media without antibiotics for several hours and plated on BHIS with 75 ng/ml anhydrotetracycline to select for double cross-out recombinants. Mutants were identified by PCR.

Insertional constructs were created by cloning into the BamHI site of pKF54 (18), transforming into *E. coli* S17-1 λ pir, and conjugally transferring to *Bacteroides* strains with integration at the *attB2* site with subsequent selection on gentamycin/cefoxitin plates.

Cloning of genes for expression from the multi-copy plasmid pFD340 (38) was performed similarly by cloning into the BamHI site of pFD340, transforming *E. coli* S17-1 λ pir with subsequent conjugal transfer to *Bacteroides* strains with selection on gentamycin/erythromycin plates.

### SDS-PAGE and western blot analyses

Cell lysates and supernatants analyzed by western immunoblots were obtained from cultures grown overnight to stationary phase in basal medium, except for strains grown for co-cultures, and for experiments where 638RΔ*lgs*_GA3_ was cultured in medium containing 50% culture supernatant as described below. For cell lysates, bacteria were pelleted and cells from the equivalent of 10 µl of culture were loaded into each well, and 20 µl of supernatants from the same cultures were loaded. These samples were prepared in 4x NuPAGE LDS Sample Buffer (Invitrogen, Waltham, MA). Samples were boiled for 5 minutes then resolved on 12% Bolt SDS-PAGE Bis/Tris gels (Invitrogen). Contents of SDS-PAGE gels were transferred to polyvinylidene difluoride (PVDF) membranes and then blocked with TBST (100 mM Tris, 0.9% NaCl, 0.05% Tween 20, pH 7.5) containing 4% skim milk for 15 min. Antiserum to Hcp_GA3_ was previously described (11, 25) and used at 1:70 dilution. Secondary antibody (alkaline phosphatase conjugated goat α-rabbit IgG) (Invitrogen) was used at a 1:2000 dilution. The membranes were developed with BCIP/NBT (5-bromo-4-chloro-3-indolylphosphate/nitro blue tetrazolium) (Seracare Life Sciences, Milford, MA).

### Co-culture experiments

For co-culture experiments, bacteria were grown in modified basal medium (MBM). All bacterial strains were resuspended from a fresh plate into MBM at an OD_600_ of ∼0.1 and then 2 ml of 638RΔ*lgs*_GA3_ or 9343Δ*lgs*_GA3_ were mixed with 2 ml of the five strains used for co-culture as shown in Fig 3A or with medium alone. The bacteria were allowed to grow for 6 hrs anaerobically and then cell lysates and supernatants were prepared for western immunoblot as described above for monocultures. All co-cultures reached very high densities as the OD_600_ of co-cultures diluted 1:3 were between 0.761 and 0.924.

### Assays using growth media with 50% supernatants from bacterial cultures

The five strains listed in Fig 3B were grown overnight to stationary phase in MBM. Bacteria were centrifuged and the supernatant was filtered using a Whatman 0.45 µM Puradisc filter (Cytiva, Marlborough, MA). 638RΔ*lgs*_GA3_ and 638R pLEC322 (p*tetR*GA3) were grown in MBM to an OD_600_ of ∼0.4 and then an equivalent volume of the cultured supernatant from the five strains were added to each. Erythromycin was added to 638R pLEC322 cultures to ensure plasmid maintenance. These co-cultures were grown for 15 hours and then cell lysates and supernatants were processed for western immunoblot analysis.

### Antagonism assays

All bacteria were grown to an OD_600_ of ∼0.7, mixed at the indicated ratio of 5:1 or 1:1 antagonist to target. 10 µl of the mixed cultures was dotted to a BHIS plate and incubated for 20 hours under anaerobic conditions. The bacterial growth was resuspended in basal medium and 10-fold serial dilutions were plated to BHIS plates and to BHIS plates with erythromycin (will grow only the target strain).

### Structural analyses

Protein structure predictions were performed with Alphafold3 (39). 3D structural similarity analysis to calculate the root mean square deviation (RMSD) was performed as described (40).

### RNASeq analyses

For RNA sequencing, bacteria were grown in triplicate in basal medium, or in basal medium with 10 µg/ml erythromycin (638R pFD340 (pvector(mc)) and 638R pLEC322 (p*tetR*_GA3_) to OD_600_ of ∼ 0.9, pelleted in the anaerobic chamber, flash-frozen and processed at the Duchossois Family Institute Microbiome Metagenomics Facility. Nucleic acid was recovered using the Maxwell RSC instrument (Promega Corporation, Madison, WI). Following DNAse treatment and elution, samples were quantified using Qubit (Life Technologies) and integrity was assessed using a Tapestation unit (Agilent Technologies, Inc., Santa Clara, CA). Ribosomal RNA was depleted from all samples using the NEBNext rRNA depletion kit (NEB). The samples were fragmented, libraries were prepared using the Ultra Directional RNA library prep kit for Illumina (NEB), and normalized libraries of biological triplicates of all samples were sequenced on the NextSeq 1000 platform (Illumina, Inc., San Diego, CA) at 2 × 100 bp read length.

Adapter and quality trimming of all reads was performed using the utilities included in the BBMap package of bioinformatics tools (v. 38.90), and the reads were mapped to the *B. fragilis* 638R genome using Bowtie 2 short-read aligner (v. 2.4.5) (41). The Bowtie 2 output was converted to sorted and indexed BAM files using SAMtools (v. 1.11) (42), and BEDtools (v. 2.30.0) (43) and compared to a General Feature Format (GFF) file containing the intervals of protein-coding domains from *B. fragilis* 638R to which were added sRNA genes detected using cmscan from infernal (v. 1.1.5) (44) and the RFam (v. 15) (45) collection of profile HMMs, using the gathering thresholds included in the models and excluding tRNA and rRNA genes detected. Read mapping results were evaluated for differential gene expression using both DESeq2 (v. 1.48.1) (46) and edgeR (v. 4.6.2) (47), running under R (v. 4.5.0) (https://www.R-project.org). Volcano plots were created in Prism, version 10.4.2 (Dotmatics, Boston, MA). RNASeq data were deposited with NCBI (BioProject ID pending).

### Data Sharing

RNASeq data were deposited with NCBI (BioProject ID pending). All other data provided in the manuscript text, figures and supplemental material. All constructs and mutants created in this study are available to the scientific community upon publication.

## Acknowledgments and funding

LT is supported by an NSF Graduate Research Fellowship 2140001, MLS is supported by NIH T32 GM144292, PAR is supported by Public Health Service grants NIH R35GM149586. This work was funded by the Duchossois Family Institute and Public Health Service grant NIH R01AI093771.

## Figure legends

**Figure S1**. Supporting data for Fig 1. **A**. Loading control for the western immunoblots shown in Figure 1B showing the top portions of the same gels and reactivity of the polyclonal antiserum to irrelevant molecules of the bacteria. Blots were probed with antiserum to the structural Hcp protein (Hcp_GA3_) encoded by the first gene of the locus (BF638R_1994). **B. B**. Clustal Omega (v. 1.2.4) (48) alignment of TetR_GA3_, TetR_1770_, and TetR_2044_. **C**. Clustal Omega alignment of TetR_1770_ and TetR_2044_.

**Figure S2**. Supporting data for Fig 2. **A**. Loading control for Fig 2C as described in legend S1. **B**. Loading control for Fig 2E. **C**. Clustal Omega alignment of Lgs_GA3_ and Lgs_BdB_.

**Figure S3**. Supporting data for Fig 3. **A**. Loading control for Fig 3A as described in Figure S1 legend. **B**. Western immunoblot analysis of the culture supernatants added 1:1 with growth medium for 638RΔ*lgs*_GA3_ from Fig 3B. **C**. Loading control for Fig 3B after culture.

**Figure S4**. Supporting data for Fig 4. Antagonism assays (same as in Fig 4) plated on BHIS without antibiotics. The bacterial dilution is indicated on the right. The antagonistic and control strains are listed along the top and the target strains are shown at the bottom.

**Figure 4.**
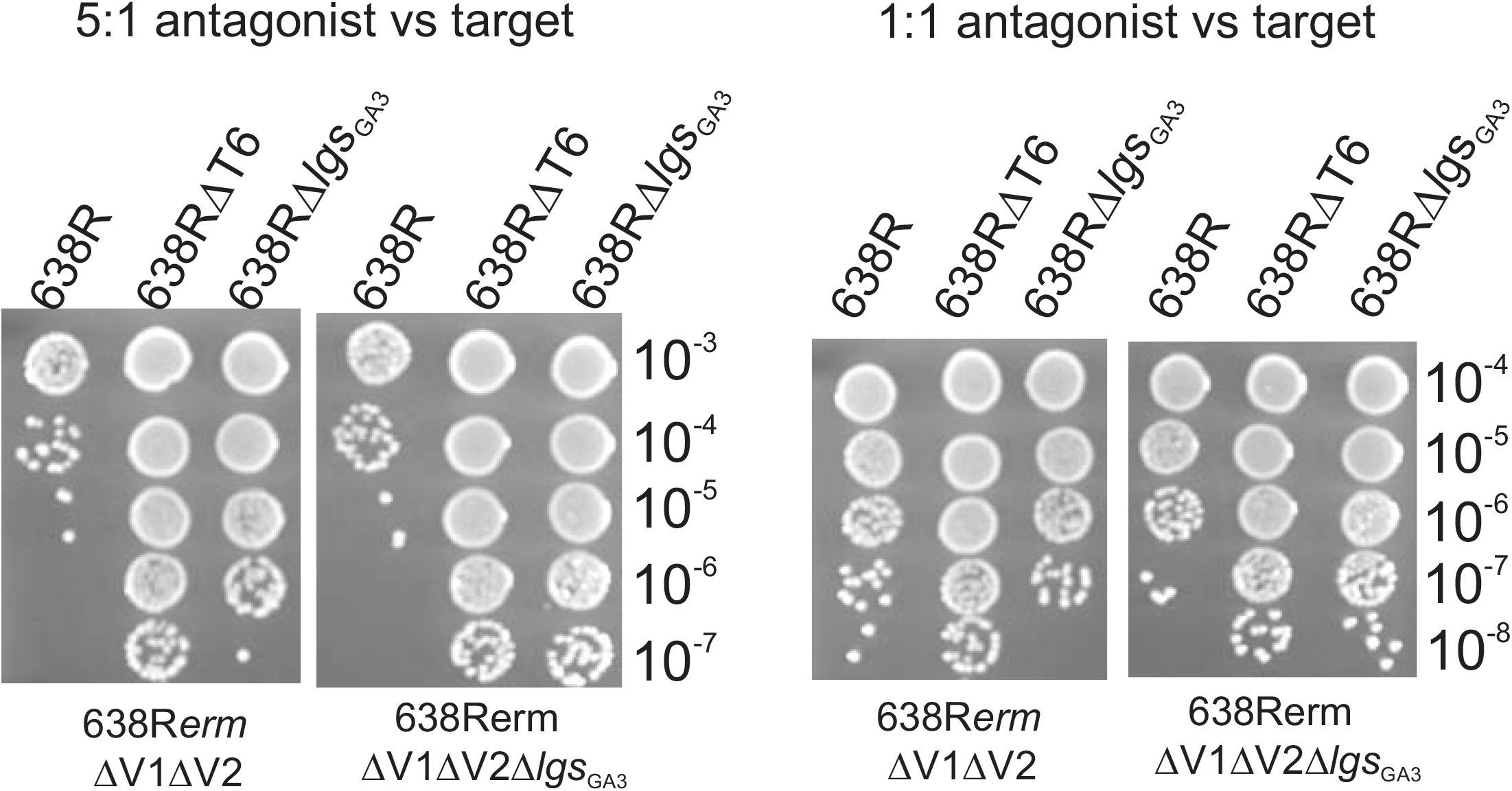
The signaling molecule produced by *lsg*_GA3_ containing target strains can induce antagonism in competing cells. Antagonism assays using isogenic strains with two different ratios (5:1 and 1:1) antagonist to target cell. Antagonistic or control strains are listed across the top and target strains are shown at the bottom. Following 20 hr co-culture, bacteria were plated in 10-fold dilutions on BHIS plates containing erythromycin, which is specific for the target strain. Fig S4 shows these same co-cultures plates on BHIS without antibiotics, allowing for quantification of each strain after co-culture.

## References

1. B. Jana, D. Salomon, Type VI secretion system: a modular toolkit for bacterial dominance. Future Microbiol 14, 1451–1463 (2019).

2. Y. Cherrak, N. Flaugnatti, E. Durand, L. Journet, E. Cascales, Structure and Activity of the Type VI Secretion System. Microbiol Spec 7 (2019).

3. L. Garcia-Bayona, L. E. Comstock, Bacterial antagonism in host-associated microbial communities. Science 361 (2018).

4. M. LeRoux, S. B. Peterson, J. D. Mougous, Bacterial danger sensing. J Mol Biol 427, 3744–3753 (2015).

5. J. T. Hespanhol, L. Nobrega-Silva, E. Bayer-Santos, Regulation of type VI secretion systems at the transcriptional, posttranscriptional and posttranslational level. Microbiology (Reading) 169 (2023).

6. R. Niehus, N. M. Oliveira, A. Li, A. G. Fletcher, K. R. Foster, The evolution of strategy in bacterial warfare via the regulation of bacteriocins and antibiotics. eLife 10 (2021).

7. D. A. I. Mavridou, D. Gonzalez, W. Kim, S. A. West, K. R. Foster, Bacteria Use Collective Behavior to Generate Diverse Combat Strategies. Curr Biol 28, 345–355 e344 (2018).

8. J. J. Faith et al., The long-term stability of the human gut microbiota. Science 341 (2013).

9. S. C. Forster et al., A human gut bacterial genome and culture collection for improved metagenomic analyses. Nat Biotechnol 37, 186–192 (2019).

10. D. T. Truong, A. Tett, E. Pasolli, C. Huttenhower, N. Segata, Microbial strain-level population structure and genetic diversity from metagenomes. Genome Res 27, 626–638 (2017).

11. M. Chatzidaki-Livanis, M. J. Coyne, L. E. Comstock, An antimicrobial protein of the gut symbiont Bacteroides fragilis with a MACPF domain of host immune proteins. Mol Microbiol 94, 1361–1374 (2014).

12. M. Chatzidaki-Livanis et al., Gut symbiont Bacteroides fragilis secretes a eukaryotic-like ubiquitin protein that mediates intraspecies antagonism. mBio 8 (2017).

13. K. G. Roelofs, M. J. Coyne, R. R. Gentyala, M. Chatzidaki-Livanis, L. E. Comstock, Bacteroidales secreted antimicrobial proteins target surface molecules necessary for gut colonization and mediate competition in vivo. mBio 7 (2016).

14. M. J. Coyne et al., A family of anti-Bacteroidales peptide toxins wide-spread in the human gut microbiota. Nat Comm 10, 3460 (2019).

15. A. M. Shumaker, V. Laclare McEneany, M. J. Coyne, P. A. Silver, L. E. Comstock, Identification of a fifth antibacterial toxin produced by a single Bacteroides fragilis strain. J Bacteriol 201 (2019).

16. J. C. Evans et al., A proteolytically activated antimicrobial toxin encoded on a mobile plasmid of Bacteroidales induces a protective response. Nat Comm 13, 4258 (2022).

17. Y. Bao et al., A common pathway for activation of host-targeting and bacteria-targeting toxins in human intestinal bacteria. mBio 12, e0065621 (2021).

18. H. L. Abrahamsen et al., Distant relatives of a eukaryotic cell-specific toxin family evolved a complement-like mechanism to kill bacteria. Nat Comm 15, 5028 (2024).

19. M. J. Coyne, K. G. Roelofs, L. E. Comstock, Type VI secretion systems of human gut Bacteroidales segregate into three genetic architectures, two of which are contained on mobile genetic elements. BMC Genomics 17, 1–21 (2016).

20. M. Basler, B. T. Ho, J. J. Mekalanos, Tit-for-tat: type VI secretion system counterattack during bacterial cell-cell interactions. Cell 152, 884–894 (2013).

21. M. L. Sheahan et al., A ubiquitous mobile genetic element changes the antagonistic weaponry of a human gut symbiont. Science 386, 414–420 (2024).

22. L. Cuthbertson, J. R. Nodwell, The TetR family of regulators. Microbiol Mol Biol Rev 77, 440–475 (2013).

23. J. Filipek et al., Comprehensive structural overview of the C-terminal ligand-binding domains of the TetR family regulators. J Struct Biol 216, 108071 (2024).

24. R. S. Patil et al., TetR and OmpR family regulators in natural product biosynthesis and resistance. Proteins 93, 38–71 (2025).

25. M. Chatzidaki-Livanis, N. Geva-Zatorsky, L. E. Comstock, Bacteroides fragilis type VI secretion systems use novel effector and immunity proteins to antagonize human gut Bacteroidales species. Proc Natl Acad Sci U S A 113, 3627–3632 (2016).

26. B. D. Ross et al., Human gut bacteria contain acquired interbacterial defence systems. Nature 575, 224–228 (2019).

27. N. Jahan et al., Insights into the biosynthesis of the Vibrio cholerae major autoinducer CAI-1 from the crystal structure of the PLP-dependent enzyme CqsA. J Mol Biol 392, 763–773 (2009).

28. R. C. Kelly et al., The Vibrio cholerae quorum-sensing autoinducer CAI-1: analysis of the biosynthetic enzyme CqsA. Nat Chem Biol 5, 891–895 (2009).

29. G. A. Hunter, G. C. Ferreira, Molecular enzymology of 5-aminolevulinate synthase, the gatekeeper of heme biosynthesis. Biochim Biophys Acta 1814, 1467–1473 (2011).

30. H. Ikushiro, H. Hayashi, H. Kagamiyama, Bacterial serine palmitoyltransferase: a water-soluble homodimeric prototype of the eukaryotic enzyme. Biochim Biophys Acta 1647, 116–120 (2003).

31. M. B. Miller, K. Skorupski, D. H. Lenz, R. K. Taylor, B. L. Bassler, Parallel quorum sensing systems converge to regulate virulence in Vibrio cholerae. Cell 110, 303–314 (2002).

32. D. A. Higgins et al., The major Vibrio cholerae autoinducer and its role in virulence factor production. Nature 450, 883–886 (2007).

33. D. Alexeev et al., The crystal structure of 8-amino-7-oxononanoate synthase: a bacterial PLP-dependent, acyl-CoA-condensing enzyme. J Mol Biol 284, 401–419 (1998).

34. S. Brameyer et al., Outer Membrane Vesicles Facilitate Trafficking of the Hydrophobic Signaling Molecule CAI-1 between Vibrio harveyi Cells. J Bacteriol 200 (2018).

35. A. Chen, Z. Jiang, M. D. Burkart, Enzymology of standalone elongating ketosynthases. Chem Sci 13, 4225–4238 (2022).

36. L. Garcia-Bayona, L. E. Comstock, Streamlined genetic manipulation of diverse Bacteroides and Parabacteroides isolates from the human gut microbiota. mBio 10 (2019).

37. A. Pantosti, A. O. Tzianabos, A. B. Onderdonk, D. L. Kasper, Immunochemical characterization of two surface polysaccharides of Bacteroides fragilis. Infect Immun 59, 2075–2082 (1991).

38. C. J. Smith, M. B. Rogers, M. L. McKee, Heterologous gene expression in Bacteroides fragilis. Plasmid 27, 141–154 (1992).

39. J. Abramson et al., Accurate structure prediction of biomolecular interactions with AlphaFold 3. Nature 630, 493–500 (2024).

40. S. Bittrich, J. Segura, J. M. Duarte, S. K. Burley, Y. Rose, RCSB protein Data Bank: exploring protein 3D similarities via comprehensive structural alignments. Bioinformatics 40 (2024).

41. B. Langmead, S. L. Salzberg, Fast gapped-read alignment with Bowtie 2. Nat Methods 9, 357–359 (2012).

42. P. Danecek et al., Twelve years of SAMtools and BCFtools. Gigascience 10 (2021).

43. A. R. Quinlan, I. M. Hall, BEDTools: a flexible suite of utilities for comparing genomic features. Bioinformatics 26, 841–842 (2010).

44. E. P. Nawrocki, S. R. Eddy, Infernal 1.1: 100-fold faster RNA homology searches. Bioinformatics 29, 2933–2935 (2013).

45. N. Ontiveros-Palacios et al., Rfam 15: RNA families database in 2025. Nucleic Acids Res 53, D258–D267 (2025).

46. M. I. Love, W. Huber, S. Anders, Moderated estimation of fold change and dispersion for RNA-seq data with DESeq2. Genome Biol 15, 550 (2014).

47. M. D. Robinson, D. J. McCarthy, G. K. Smyth, edgeR: a Bioconductor package for differential expression analysis of digital gene expression data. Bioinformatics 26, 139–140 (2010).

48. F. Madeira et al., The EMBL-EBI Job Dispatcher sequence analysis tools framework in 2024. Nucleic Acids Res 52, W521–W525 (2024).

