## Supplemental Figures and Tables for "Antagonism by the Type VI secretion system of *Bacteroides fragilis* is controlled by a TetR family regulator and released small molecule"

Fig. S1

A.

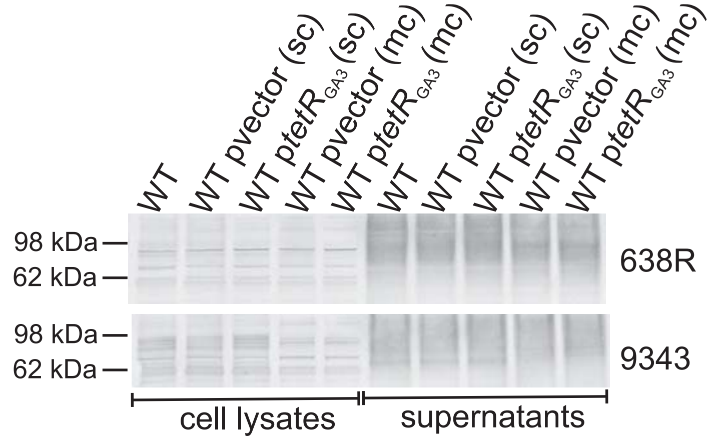

B.

```
TetRGA3         MGQGDKKIRIRRTNEQLDKEVISEFEKLVGEFGFGNVNLSALMKAADLEANVFYRRYGSM
TetR1770        MKQEEKKTRNRRTNEQLDRDVMSELEKLVAEQGFSNVNLSTLTKAAGMEANVFYRRYGSM
TetR2044        MKQEEKKNRNRRTNEQLDRDVMSELEKLVAEQGFGNVNLSTLTKAAGMEANVFYRRYGSM
                * * :** * ********::*:**:****.* **.*****:* ***.:************

TetRGA3         DNLYDRLAKQYDFWINNTIDISTLNTLGPKKFFAETFKTLFRNLSENSVMQKLLLYEMTT
TetR1770        DNLYDRLAKQYDFWMNNAIDISNLNALGPKKFFAETFKTLFRNLSDNPVMQKLLLYEMSA
TetR2044        DNLYDRLAKQYDFWMNNAIDISNLNVLGPKKFFAETFKTLFRNLSDNPVMQKLLLYEMSV
                **************:**:****.**.*******************:* **********:.

TetRGA3         INSTTKRSAETRDVMNLNLITFYENLFAPAKINIKSIASILIGGIYYLILHKECAKICTI
TetR1770        VNDTTKRTAETRDIMNLNLITYYETLFKPAKVNIKSIAAILIGGIYYLILHKECAKICTI
TetR2044        VNDTTKRTAETRDIMNLNLITYYETLFKPAKVNIKSIAAILIGGIYYLILHKECAKICTI
                :*.****:*****:*******:**.** ***:******:*********************

TetRGA3         DYKTKEGENAFSEGIDFLTDIIFDRLEMYDRDKKAIRQMISDGISESKICKYMGINKNDL
TetR1770        DFNTPEGEKAFSDGIDFLTDTIFNRLEAHERDRNAVRQMLADGISELKICKYMGISKNDL
TetR2044        DFNTPEGEKAFSEGIDFLTDTIFNRLEAYERDRNAVRQMLADGISELKICKYMGISKNDL
                *::* ***:***:******* **:*** ::**::*:***::***** ********.****

TetRGA3         KTLLSE
TetR1770        KMLLSK
TetR2044        KVLLSK
                * ***:
```

C.

```
TetR1770        MKQEEKKTRNRRTNEQLDRDVMSELEKLVAEQGFSNVNLSTLTKAAGMEANVFYRRYGSM
TetR2044        MKQEEKKNRNRRTNEQLDRDVMSELEKLVAEQGFGNVNLSTLTKAAGMEANVFYRRYGSM
                *******.**************************.*************************

TetR1770        DNLYDRLAKQYDFWMNNAIDISNLNALGPKKFFAETFKTLFRNLSDNPVMQKLLLYEMSA
TetR2044        DNLYDRLAKQYDFWMNNAIDISNLNVLGPKKFFAETFKTLFRNLSDNPVMQKLLLYEMSV
                *************************.*********************************.

TetR1770        VNDTTKRTAETRDIMNLNLITYYETLFKPAKVNIKSIAAILIGGIYYLILHKECAKICTI
TetR2044        VNDTTKRTAETRDIMNLNLITYYETLFKPAKVNIKSIAAILIGGIYYLILHKECAKICTI
                ************************************************************

TetR1770        DFNTPEGEKAFSDGIDFLTDTIFNRLEAHERDRNAVRQMLADGISELKICKYMGISKNDL
TetR2044        DFNTPEGEKAFSEGIDFLTDTIFNRLEAYERDRNAVRQMLADGISELKICKYMGISKNDL
                ************:***************:*******************************

TetR1770        KMLLSK
TetR2044        KVLLSK
                *:****
```

Fig. S2

A.

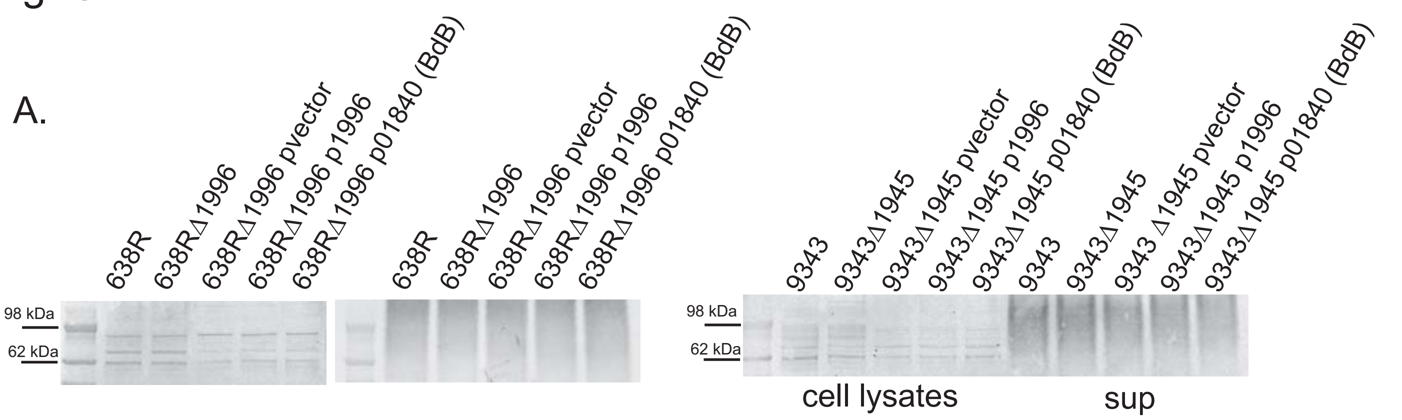

B.

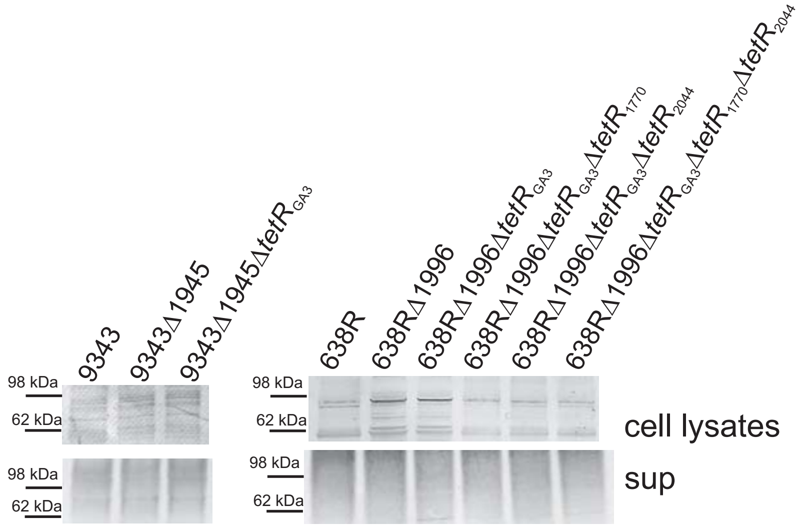

C.

```
LgsGA3   RFKIDYSVCKNSPGGSILEKVSNFQKELQLHEENSLHIYVKSPMVSGCGREVEVVDRKTN  66
         R+ I+ +  K S  GSIL KV +F+ E+ LHE++ L++Y K P ++ C REV + +R T
LgsBdB   RYVINITKNKTSKDGSILGKVLDFKNEMTLHEQSELNVYKKHPQLTACNREVTIYNRYTQ  61

         EVKKMLMFGSNSYLDATGIPSVVEKAVRVITDYGVGSGGVPLLSGTTIFQNELEKEIAKL  126
         +   + +MFGSNSYL AT   P   ++KA+ V   + G+GSGGVPLLSGT ++Q ELE  IAK+
         QKVQTIMFGSNSYLGATIFPEAIQKAIEVTKELGIGSGGVPLLSGTFMYQQELEHLIAKM  121

         TGFDDTILFSSGFTANIGVIVGLIRPNNLLVYDRLNHASLIDGALMSGAKMVRYKHNDPK  186
         +G DD+ILFSSG+TAN+GVI GLIR NNL+++D+LNHASL+DG +MSGAKM+RYKH D
         SGMDDSILFSSGYTANLGVISGLIRANNLIIHDKLNHASLLDGTVMSGAKMIRYKHGDIS  181

         ALEKILKENAGQYKDGMMVVTDGVFSMDGDIADIPAILEITKKYNALLLIDDAHATGVIG  246
          LE +LK+N+ +Y +G++VVTDGVFSMDGDIA +P ILE  +KYNA+LLID+AHATGVIG
         HLESLLKDNSPKYPEGILVVTDGVFSMDGDIAKLPLILETIRKYNAVLLIDEAHATGVIG  241

         EDGAGTLSYYDIKERENIIVTGTLSKAIGSIGGFITAKQNIIDYLRVYARSNMYSTSLPQ  306
         ++GAGTLS+Y I +R+NII+TGTLSKAIG++GG+ITA Q+IIDYLR+YARSNM+STSLP
         DNGAGTLSHYGITDRKNIILTGTLSKAIGTVGGYITASQDIIDYLRIYARSNMFSTSLPP  301

         SICAASLEVIKEMRNTDIQNALKRNAEYVRNGLKALGFNTLNSMTPIIPVIVGDEYILTQ  366
         S+CAA++EVIK M+NT+I   LK+NA+Y+++  +  GFN LN+ TPIIP++VGD+ ILTQ
         SVCAAAIEVIKVMQNTNIVERLKQNAKYLQHKFRDGGFNILNTETPIIPLVVGDQAILTQ  361

         ITKELYDRDIFTNAIFPPVVPPNMCRIRIGVMSSHTFEDCDRLINVFHEIGKKY  420
         + K+ +D     N+IFPPVVP N+ R RI +M++HT +D D L  +  ++  KY
         MAKDAFDNGFVLNSIFPPVVPANLTRFRISIMANHTQKDMDMLFELMIKLFDKY  415
```

Fig. S3

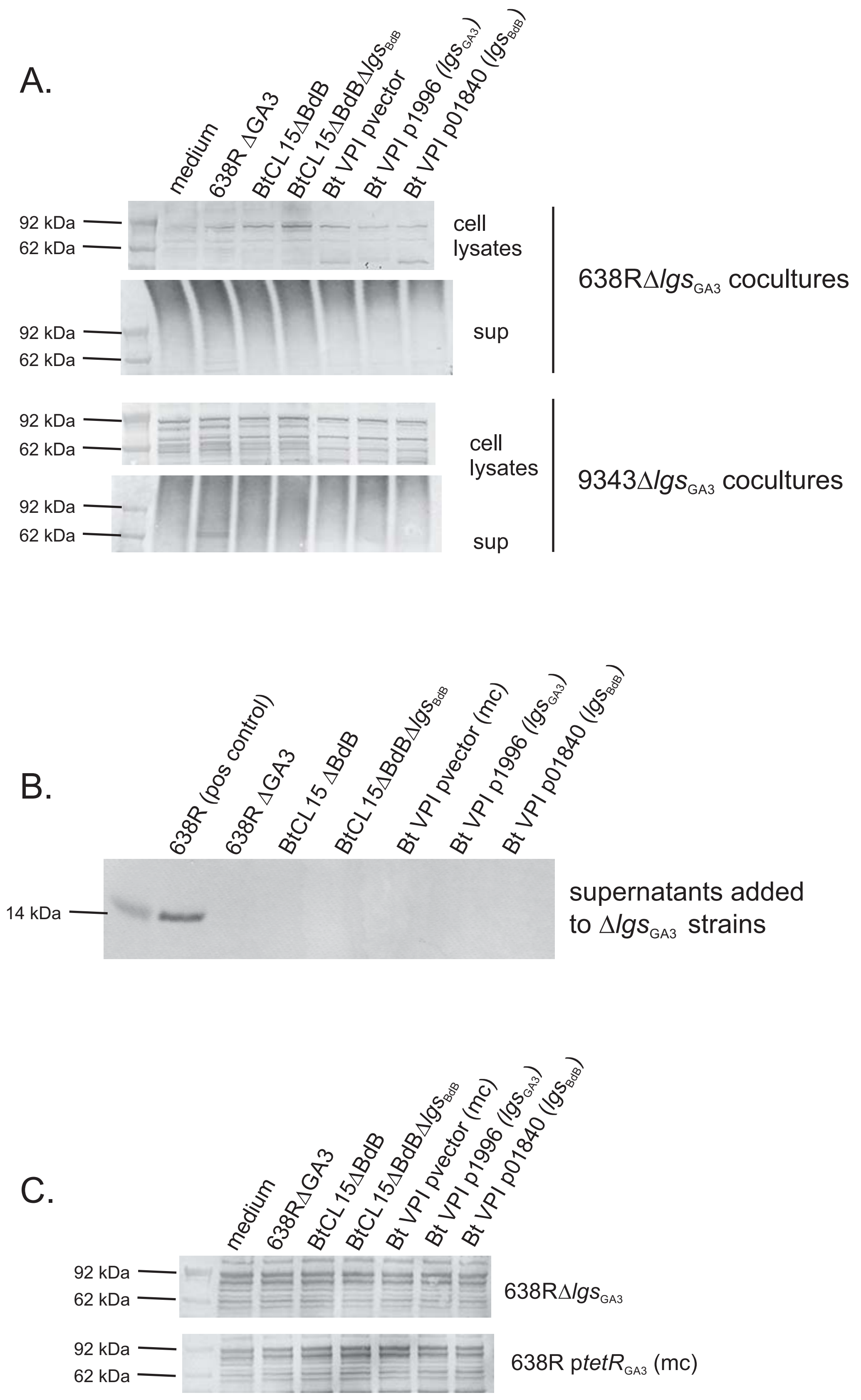
A.
medium
638R ΔGA3
BtCL15ΔBdB
BtCL15ΔBdBΔlgsBdB
Bt VPI pvector
Bt VPI p1996 (lgsGA3)
Bt VPI p01840 (lgsBdB)
92 kDa
62 kDa
cell lysates
sup
638RΔlgsGA3 cocultures
9343ΔlgsGA3 cocultures
B.
638R (pos control)
638R ΔGA3
BtCL15 ΔBdB
BtCL15ΔBdBΔlgsBdB
Bt VPI pvector (mc)
Bt VPI p1996 (lgsGA3)
Bt VPI p01840 (lgsBdB)
14 kDa
supernatants added to ΔlgsGA3 strains
C.
medium
638RΔGA3
BtCL15ΔBdB
BtCL15ΔBdBΔlgsBdB
Bt VPI pvector (mc)
Bt VPI p1996 (lgsGA3)
Bt VPI p01840 (lgsBdB)
92 kDa
62 kDa
638RΔlgsGA3
638R ptetRGA3 (mc)

Fig. S4

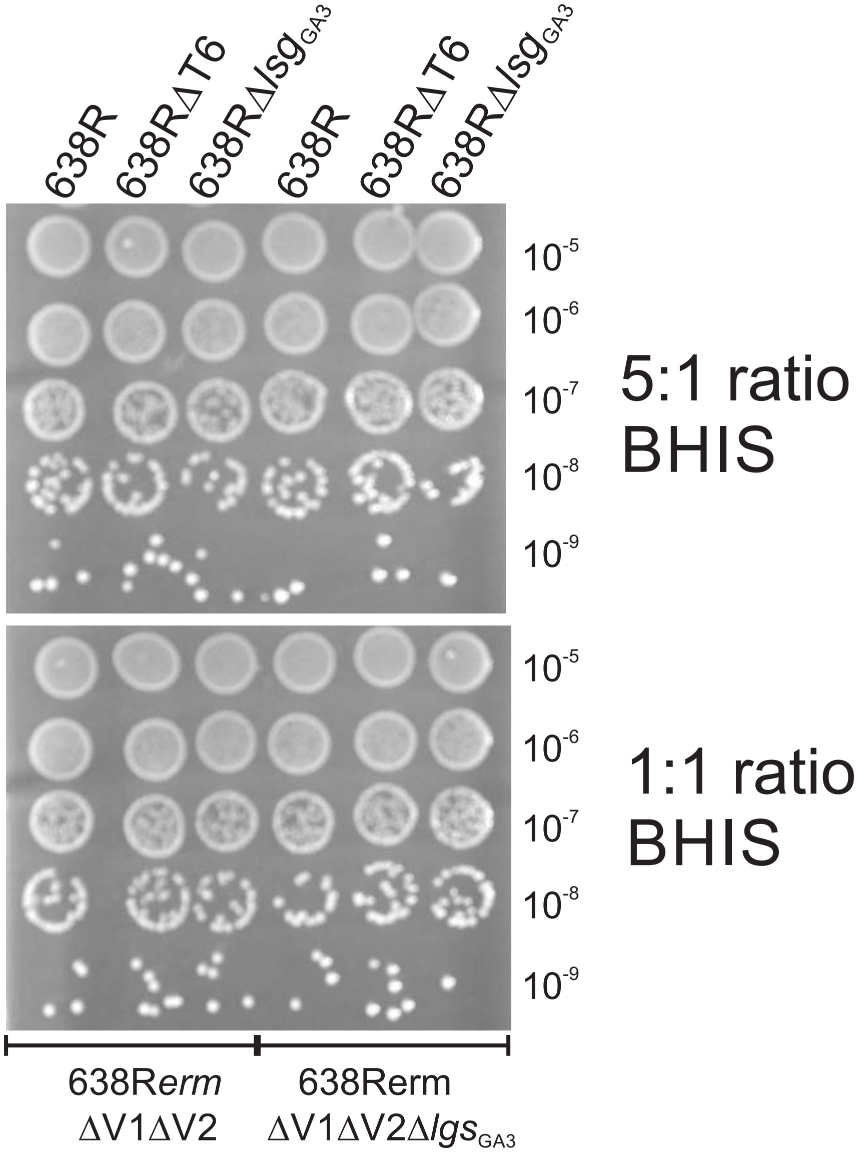
638R
638RΔT6
638RΔ$lsg_{GA3}$
638R
638RΔT6
638RΔ$lsg_{GA3}$
$10^{-5}$
$10^{-6}$
$10^{-7}$
$10^{-8}$
$10^{-9}$
5:1 ratio
BHIS
$10^{-5}$
$10^{-6}$
$10^{-7}$
$10^{-8}$
$10^{-9}$
1:1 ratio
BHIS
638R*erm*
ΔV1ΔV2
638Rerm
ΔV1ΔV2Δ$lgs_{GA3}$

**Table S1. Primers used in this study**

| | | |
|---|---|---|
| Clone $tetR_{GA3}$ ((BF638R_1995) into pFD340 | forward | tgacggatccatacctccggacttctggtg |
| Resulting plasmid pLEC322 (a.k.a p$tetR_{GA3}$ (mc)) | reverse | tgacggatcccatctgtccatatagtctcctgtgac |
| Clone $tetR_{GA3}$ into pKF54 | forward | aatcagaattgactctagagaataagtactatataataatgatgggac |
| Resulting plasmid pLEC497 (a.k.a. p$tetR_{GA3}$ (sc)) | reverse | tcgaattcctgcagcccggggtctcctgtgactattcc |
| Delete BF638R_1996 ($lgs_{GA3}$) | left flank forward | ggcatagtatcagatgagtggatttccttaaacatctgtcc |
| Resulting plasmid pLEC494 | left flank reverse | ctatttcatggaacctagatatgttctctttc |
| | right flank forward | atctaggttccatgaaataggaaaaaaatacgg |
| | right flank reverse | cgaattcctgcagcccggggatagagcccgtagttacg |
| Delete BF638R_1995-1996 ($tetR_{GA3}$,$lgs_{GA3}$) - made in pMLS36 | left flank forward | ggcatagtatcagatgagtgcacagatacagacttctc |
| Resulting plasmid pLEC500 | left flank reverse | cctgtgactaggacttatctgataaaacgg |
| | right flank forward | agataagtcctagtcacaggagactatatg |
| | right flank reverse | cgaattcctgcagcccggggctgttcaactatgagggag |
| Delete 638R $tetR_{1770}$ - Construct made in pMLS36 | Left flank forward | ggcatagtatcagatgagtgcaattaaagccggaacag |
| Resulting plasmid pLEC504 | Left flank reverse | gccgtggttactcttcctgcttcattcttatatc |
| | Right flank forward | gcaggaagagtaaccacggcacaactttttatatag |
| | Right flank reverse | cgaattcctgcagcccgggggccaaaacattacctgcttg |
| Delete 638R $tetR_{2044}$ - Construct made in pMLS36 | Left flank forward | ggcatagtatcagatgagtgggaagctgttaatcttgtc |
| Resulting plasmid pLEC505 | Left flank reverse | agagcaacactcttatatcatatttataccggc |
| | Right flank forward | tgatataagagtgttgctctcaaagtaac |
| | Right flank reverse | cgaattcctgcagcccgggggtccaatacttccaacaatg |
| Clone BF638R_1996 ($lgs_{GA3}$) into pFD340 | forward | aatcagaattgactctagaggcaaacaaatttataacgtataaatac |
| Resulting plasmid pLEC491 | reverse | attcgagctcggtacccggggttttatcagataagtccgtattttttc |
| Clone $lgs_{GA3}$ into pKF54 | forward | aatcagaattgactctagagaacgtataaatactaaaatgaaagag |
| Resulting plasmid pLEC493 | reverse | tcgaattcctgcagcccgggtatgccgttttatcagataag |
| Clone Bd-B ligand synthase ($lgs_{BdB}$ EH213_01840) into pFD340 | forward | aatcagaattgactctagagcacagtctatactgtataatacataac |

| | | |
|---|---|---|
| Resulting plasmid pLEC492 | reverse | attcgagctcggtacccggggaatgattatctcacgaatcg |
| Delete BtCL15C11 *lgs*$_{BdB}$ – construct made in pLGB13 | left flank forward | taagattagcattatgagtggttataccttatatataacaatagcaac |
| Resulting plasmid pLEC503 | left flank reverse | agaatgattaaacgtacctcataatgttatg |
| | right flank forward | gaggtacgtttaatcattctataagttgaggtg |
| | right flank reverse | cgaattcctgcagcccggggctgaaggaagcttttaagc |
| Delete BF9343_1945 (*lgs*$_{GA3}$) | left flank forward | ggcatagtatcagatgagtgcttattgatgcccatatacttg |
| Resulting plasmid pTEP5 | left flank reverse | tgccgttttatttagtatttatacgttataaatttgtttg |
| | right flank forward | aaatactaaataaaacggcatattgtcatac |
| | right flank reverse | cgaattcctgcagcccggggggataaaacacagatacagac |
| Delete BF9343_1944-1945 (*tetR*$_{GA3,}$*lgs*$_{GA3}$) | left flank forward | ggcatagtatcagatgagtgggcttcggaagtatcaac |
| Resulting plasmid pTEP6 | left flank reverse | tgccgttttatagtcacaggagactatatg |
| | right flank forward | cctgtgactataaaacggcatattgtcatacaaataatag |
| | right flank reverse | cgaattcctgcagcccggggccgttcagtctatggccg |

**Table S2. Strains and plasmids used in this study**

| **Bacterial strains** | **source** |
|---|---|
| *Bacteroides fragilis* 638R | lab stock |
| *Bacteroides fragilis* NCTC9343 | lab stock |
| *Bacteroides thetaiotaomicron* VPI 8452 | lab stock |
| *Bacteroides thetaiotaomicron* CL15T12C11 $\Delta$Bd-B (EH213_01844) | (1) |
| *Bacteroides thetaiotaomicron* CL15T12C11 $\Delta$Bd-B $\Delta lgs_{BdB}$ (EH213_01840) | this study |
| *Bacteroides fragilis* 638R $\Delta tetR_{GA3}$ (Bf638R_1995) | this study |
| *Bacteroides fragilis* 638R $\Delta tetR_{GA3}$ $\Delta tetR_{1770}$ (BF638R_1770) | this study |
| *Bacteroides fragilis* 638R $tetR_{GA3}\Delta tetR_{2044}$ (BF638R_2044) | this study |
| *Bacteroides fragilis* 638R $\Delta tetR_{GA3}$ $\Delta tetR_{1770}$ $\Delta tetR_{2044}$ | this study |
| *Bacteroides fragilis* 638R pLEC497 | this study |
| *Bacteroides fragilis* 638R pLEC322 | this study |
| *Bacteroides fragilis* 9343 pLEC497 | this study |
| *Bacteroides fragilis* 9343 pLEC322 | this study |
| *Bacteroides fragilis* 638R $\Delta lgs_{GA3}$ (Bf638R_1996) | this study |
| *Bacteroides fragilis* 638R $\Delta tetR_{GA3}$ $\Delta lgs_{GA3}$ | this study |
| *Bacteroides fragilis* 638R $\Delta tetR_{GA3}$ $\Delta lgs_{GA3}$ $\Delta tetR_{1770}$ | this study |
| *Bacteroides fragilis* 638R $\Delta tetR_{GA3}$ $\Delta lgs_{GA3}$ $\Delta tetR_{2044}$ | this study |
| *Bacteroides fragilis* 638R $\Delta tetR_{GA3}$ $\Delta lgs_{GA3}$ $\Delta tetR_{1770}$ $\Delta tetR_{2044}$ | this study |
| *Bacteroides fragilis* 9343 $\Delta lgs_{GA3}$ (Bf9343_1945) | this study |
| *Bacteroides fragilis* 9343 $\Delta tetR_{GA3}$ $\Delta lgs_{GA3}$ | this study |
| *B. fragilis* 638R $\Delta$T6 ($\Delta$T6 is deletion of BF638R_1991-1993) | (2) |
| *B. fragilis* 638R $\Delta$T6 $\Delta lgs_{GA3}$ | this study |
| *B. fragilis* 638R $\Delta$GA3 ($\Delta$638R_1994-1971) | (3) |
| *B. fragilis* 638R *ermG* $\Delta$V1$\Delta$V2 | (3) |
| *B. fragilis* 638R *ermG* $\Delta$V1$\Delta$V2 $\Delta lgs_{GA3}$ | this study |
| *Bacteroides fragilis* 638R $\Delta lgs_{GA3}$ pLEC491 | this study |
| *Bacteroides fragilis* 638R $\Delta lgs_{GA3}$ pLEC492 | this study |
| *Bacteroides fragilis* 638R $\Delta lgs_{GA3}$ pLEC493 | this study |
| *Bacteroides fragilis* 638R $\Delta lgs_{GA3}$ pLEC501 | this study |
| *Bacteroides fragilis* 638R $\Delta$Bf638R_1995 pFD340 (vector control) | this study |
| *Bacteroides fragilis* 638R $\Delta$Bf638R_1995 pLEC322 | this study |
| *Bacteroides fragilis* 638R $\Delta$Bf638R_1995 pKF54 (vector control) | this study |
| *Bacteroides fragilis* 638R $\Delta$Bf638R_1995 pLEC497 | this study |
| *Bacteroides thetaiotaomicron* VPI 8452 pFD340 (vector control) | this study |
| *Bacteroides thetaiotaomicron* VPI 8452 pLEC491 | this study |
| *Bacteroides thetaiotaomicron* VPI 8452 pLEC492 | this study |
| *E. coli* S17-1 $\lambda$ *pir* | E. Martens |
| **Plasmids** | |
| pMLS36 – suicide vector for mutant construction | (3) |
| pFD340 – expression plasmid (a.k.a. pvector (mc)) | (4) |
| pKF54 – integrative expression plasmid (a.k.a. pvector (sc)) | (5) |
| pLGB13 – vector for creation of mutations with counterselection | (6) |

| | |
|---|---|
| pLEC322 – BF638R_1995 (*tetR*$_{GA3}$) cloned into the BamHI site of pFD340 (a.k.a p*tetR*$_{GA3}$ (mc)) | This study |
| pLEC491 - 638R_1996 gene cloned into BamHI site of pFD340 (a.k.a. p1996) | this study |
| pLEC492 - BtCL15 gene EH213_01840 of Bd-B region cloned into BamHI site of pFD340 (p01840) | this study |
| pLEC494 - To delete Bf638R_1996. Flanks cloned into BamHI site of pMLS36 | this study |
| pLEC497 – *tetR*$_{GA3}$ cloned into BamHI site of pKF54 (a.k.a. p*tetR*$_{GA3}$ (sc)) | this study |
| pLEC500 - To delete BF638R_1995-1996. Flanks cloned into the BamHI site of pMLS36 | this study |
| pLEC503 - To delete BdB ligand synthase of BtCL15. Flanks cloned into BamHI pLGB13 | this study |
| pLEC504 – To delete Bf638R_1770 – cloned into BamHI site pMLS36 | this study |
| pLEC505 – To delete Bf638R_2044 – cloned into BamHI site pMLS36 | this study |
| pTEP5 – To delete *tetR*$_{GA3}$ of 9343 (BF9343_1944) – cloned into BamHI site of pMLS36 | this study |
| pTEP6 – To delete *tetR*$_{GA3}$ and *lgs*$_{GA3}$ (BF9343_1944-1945) – cloned into BamHI site of pMLS36 | this study |

1. M. J. Coyne *et al.*, A family of anti-Bacteroidales peptide toxins wide-spread in the human gut microbiota. *Nat Comm* **10**, 3460 (2019).
2. M. Chatzidaki-Livanis, N. Geva-Zatorsky, L. E. Comstock, *Bacteroides fragilis* type VI secretion systems use novel effector and immunity proteins to antagonize human gut Bacteroidales species. *Proc Natl Acad Sci U S A* **113**, 3627-3632 (2016).
3. M. L. Sheahan *et al.*, A ubiquitous mobile genetic element changes the antagonistic weaponry of a human gut symbiont. *Science* **386**, 414-420 (2024).
4. C. J. Smith, M. B. Rogers, M. L. McKee, Heterologous gene expression in *Bacteroides fragilis*. *Plasmid* **27**, 141-154 (1992).
5. H. L. Abrahamsen *et al.*, Distant relatives of a eukaryotic cell-specific toxin family evolved a complement-like mechanism to kill bacteria. *Nat Comm* **15**, 5028 (2024).
6. L. Garcia-Bayona, L. E. Comstock, Streamlined genetic manipulation of diverse *Bacteroides* and *Parabacteroides* isolates from the human gut microbiota. *mBio* **10** (2019).
